# A toxic friend: Genotoxic and mutagenic activity of the probiotic strain *Escherichia coli* Nissle 1917

**DOI:** 10.1101/2021.03.22.436450

**Authors:** Jean-Philippe Nougayrède, Camille Chagneau, Jean-Paul Motta, Nadège Bossuet-Greif, Marcy Belloy, Frédéric Taieb, Jean-Jacques Gratadoux, Muriel Thomas, Philippe Langella, Eric Oswald

## Abstract

The probiotic *Escherichia coli* strain Nissle 1917 (DSM 6601, Mutaflor), generally considered as beneficial and safe, has been used for a century to treat various intestinal diseases. However, Nissle 1917 hosts in its genome the *pks* pathogenicity island that codes for the biosynthesis of the genotoxin colibactin. Colibactin is a potent DNA alkylator, suspected to play a role in colorectal cancer development. We show in this study that Nissle 1917 is functionally capable of producing colibactin and inducing interstrand crosslinks in the genomic DNA of epithelial cells exposed to the probiotic. This toxicity was even exacerbated with lower doses of the probiotic, when the exposed cells started to divide again but exhibited aberrant anaphases and increased gene mutation frequency. DNA damage was confirmed *in vivo* in mouse models of intestinal colonization, demonstrating that Nissle 1917 produces the genotoxin in the gut lumen. Although it is possible that daily treatment of adult humans with their microbiota does not produce the same effects, administration of Nissle 1917 as a probiotic or as a chassis to deliver therapeutics might exert long term adverse effects and thus should be considered in a risk versus benefit evaluation.

**Importance:** Nissle 1917 is sold as a probiotic and considered safe even though it is known since 2006 that it encodes the genes for colibactin synthesis. Colibactin is a potent genotoxin that is now linked to causative mutations found in human colorectal cancer. Many papers concerning the use of this strain in clinical applications ignore or elude this fact, or misleadingly suggest that Nissle 1917 does not induce DNA damage. Here, we demonstrate that Nissle 1917 produces colibactin *in vitro* and *in vivo* and induces mutagenic DNA damage. This is a serious safety concern that must not be ignored, for the interests of patients, the general public, health care professionals and ethical probiotic manufacturers.

## Introduction

*Escherichia coli* Nissle 1917 is an intestinal strain originally isolated during the first world war. Nissle 1917 is a potent competitor of different enteropathogens in the gut (Nissle, 1959). Consequently, it has been used for a century as a treatment for diarrhea and more recently for other intestinal disorders such as inflammatory bowel diseases (IBDs). The use of Nissle 1917 is recommended for maintaining remission in ulcerative colitis (Floch et al., 2011; Kruis et al., 2004). It is used as a probiotic in human medicine in Germany, Australia, Canada and other countries under the name of “Mutaflor”. Nissle 1917 is also a popular chassis to engineer therapeutic bacteria for vaccine, diagnostics, biosensors and drug development (Ou et al., 2016). The popularity of Nissle 1917 resides not only in its “natural” beneficial properties, but also in the general acceptance that it is harmless and safe. Its safety profile is based in part on the belief that Nissle 1917 does not produce any toxin associated with pathogenic strains of *E. coli*. Although this statement is still propagated in the recent biomedical literature, it was shown in 2006 that Nissle 1917 hosts a 54 kb *pks* island coding for non-ribosomal and polyketide synthases (NRPS and PKS) allowing synthesis of a hybrid peptide-polyketide metabolite called colibactin (Homburg et al., 2007; Nougayrède et al., 2006).

Colibactin is a genotoxin that binds and crosslinks the opposite strands of DNA, resulting in DNA damages and gene mutagenesis in eukaryotic cells (Bossuet-Greif et al., 2018; Cuevas-Ramos et al., 2010; Dziubańska-Kusibab et al., 2020; Iftekhar et al., 2021; Nougayrède et al., 2006; Pleguezuelos-Manzano et al., 2020; Wilson et al., 2019). Colibactin is a virulence factor during systemic infection (Marcq et al., 2014; Martin et al., 2013; McCarthy et al., 2015), and plays a substantial role in colorectal cancer. Indeed, colibactin-producing *E. coli* promote colorectal cancer in mouse models (Arthur et al., 2012; Cougnoux et al., 2014) and the DNA mutational signature of colibactin has been found in cohorts of patients with colorectal cancer, including in the APC cancer driver gene (Dziubańska-Kusibab et al., 2020; Pleguezuelos-Manzano et al., 2020; Terlouw et al., 2020). A conflicting report claimed that “no genotoxicity is detectable for *E. coli* strain Nissle 1917 by standard *in vitro* and *in vivo* tests” (Dubbert et al., 2020) but the authors used assays that are suboptimal to demonstrate production and mutagenicity of colibactin, such as the use of *Salmonella* reporter bacteria that are killed by the microcins produced by Nissle 1917 (Massip et al., 2020; Sassone-Corsi et al., 2016). Recently, in a study using stem cell-derived human intestinal organoids to evaluate the safety of the probiotic, Nissle 1917 “was found to be safe” (Pradhan and Weiss, 2020), while exposure of such organoids to *pks*+ *E. coli* induced the colibactin-specific mutational signature (Pleguezuelos-Manzano et al., 2020). Here, we examined the production and genotoxicity of colibactin by Nissle 1917 *in vitro*, using assays adapted to the described mode of action of the toxin, and *in vivo* in two mouse models.

## Results

### Nissle 1917 produces colibactin and induces DNA crosslinks in infected epithelial cells

DNA interstrand crosslinks generated by colibactin impair the denaturation of DNA and thus inhibits its electrophoretic mobility in denaturing conditions (Bossuet-Greif et al., 2018). We examined whether infection of epithelial cells with Nissle 1917 could induce crosslinks in the host genomic DNA. Cultured human epithelial HeLa cells were exposed to live *E. coli* Nissle 1917 for 4 hours, then the cell genomic DNA was purified and analyzed by denaturing gel electrophoresis. In contrast to the DNA of control cells which migrated as a high molecular weight band, a fraction of the DNA of the cells exposed to Nissle 1917 remained in the loading well (Fig 1). Similar genomic DNA with impaired electrophoretic migration was observed in cells treated with cisplatin, a DNA crosslinking agent (Fig 1a). In contrast, a Nissle 1917 mutant for the phosphopantetheinyl transferase ClbA, required for activation of the NRPS and PKS in the *pks* pathway (Martin et al., 2013), did not induce non-migrating genomic DNA (Fig 1ab). Similarly, no crosslinking activity was detected with the Nissle 1917 strain mutated for the peptidase ClbP that cleaves the inactive precolibactin to generate the mature active colibactin (Brotherton and Balskus, 2013) (Fig 1 ab). We also observed the DNA crosslinking activity in exogenous DNA exposed to the wild-type Nissle 1917 but not to the *clbA* and *clbP* mutants (Sup Fig 1). Thus, Nissle 1917 synthesizes mature DNA crosslinking colibactin.

**Figure 1.**
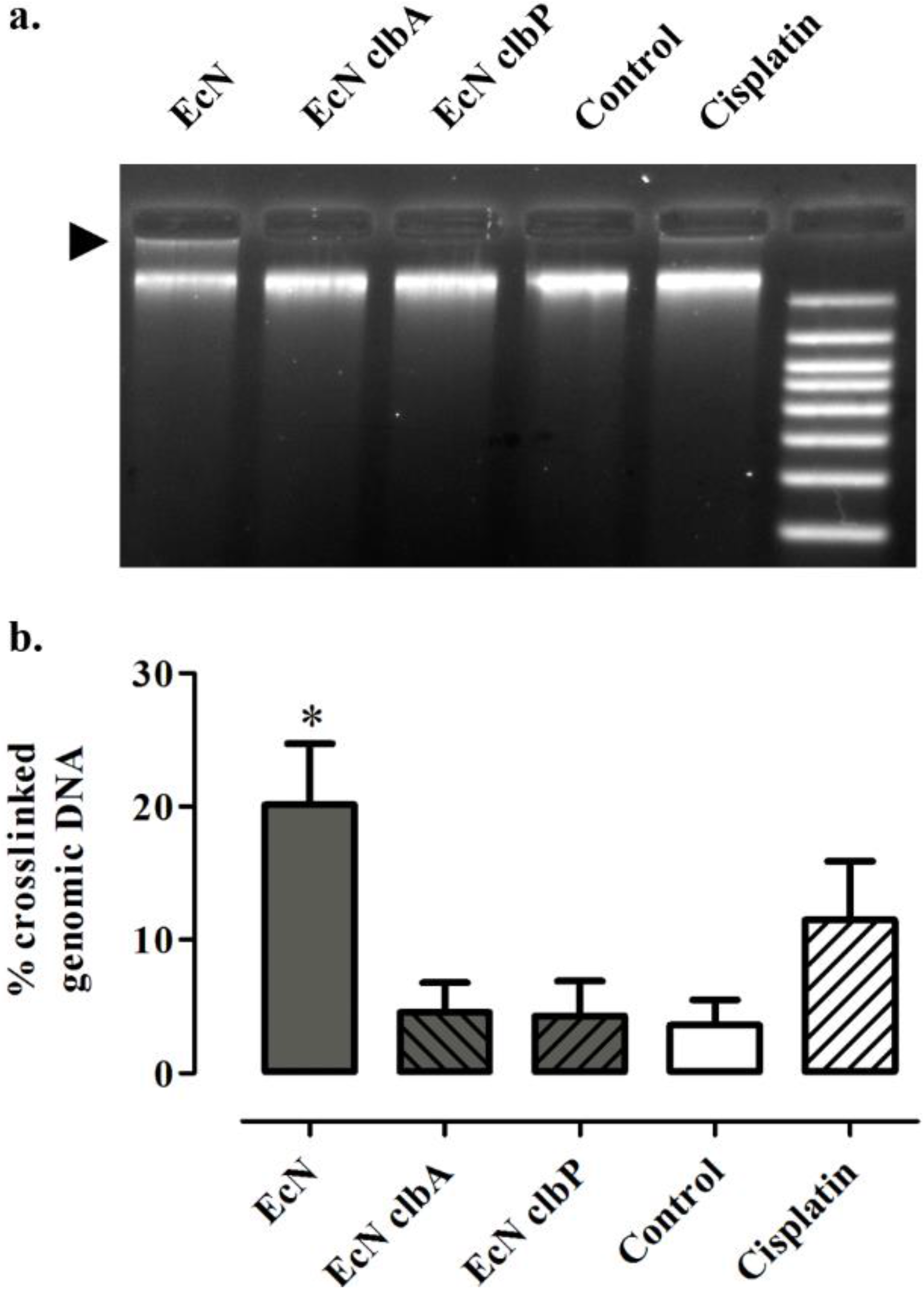
*E. coli* Nissle 1917 produces colibactin and induces interstrand crosslinks in the host cell genomic DNA. (a) HeLa cells were infected for 4 h, at a multiplicity of infection of 400 bacteria per cell, with *E. coli* Nissle (EcN), *clbA* or *clbP* isogenic mutants, left uninfected or treated 4 h with 100 μM cisplatin. Then, the cell genomic DNA was purified and analyzed by denaturing electrophoresis. The arrow points to the non-migrating DNA that remains in the loading well. (b) The DNA signal in the upper non-migrating band relative to the total DNA signal in the lane was determined by image analysis in ImageJ. The mean % of crosslinked DNA and standard error of the mean (n=3 independent experiments) are shown. * p<0.05 compared to control, one-way ANOVA with Dunnett posttest compared to control.

### Infection with Nissle 1917 induces the recruitment of the DNA repair machinery

It was recently shown that upon formation of DNA crosslinks by colibactin, the cells recruit the kinase ataxia telangiectasia and Rad3-related (ATR), which phosphorylate the Ser33 of the replication protein A-32 (RPA) in nuclear DNA repair foci together with phosphorylated histone γH2AX (Bossuet-Greif et al., 2018). Immunofluorescence of Ser33-phosphorylated RPA and γH2AX showed nuclear foci of both markers in HeLa cells 4 h after infection with Nissle 1917, or following treatment with the crosslinking drug cisplatin, but not after infection with the *clbA* or *clbP* mutants (Fig 2a). The γH2AX and p-RPA foci increased with the multiplicity of infection (MOI) with the wild-type Nissle 1917, and remained plainly measurable 20 h after infection, even at the low MOI of 20 bacteria per cell (Fig 2b). Together these results demonstrate that Nissle 1917 induces dose and time dependent DNA crosslinks in exposed cells, resulting in cognate DNA repair machinery recruitment.

**Figure 2.**
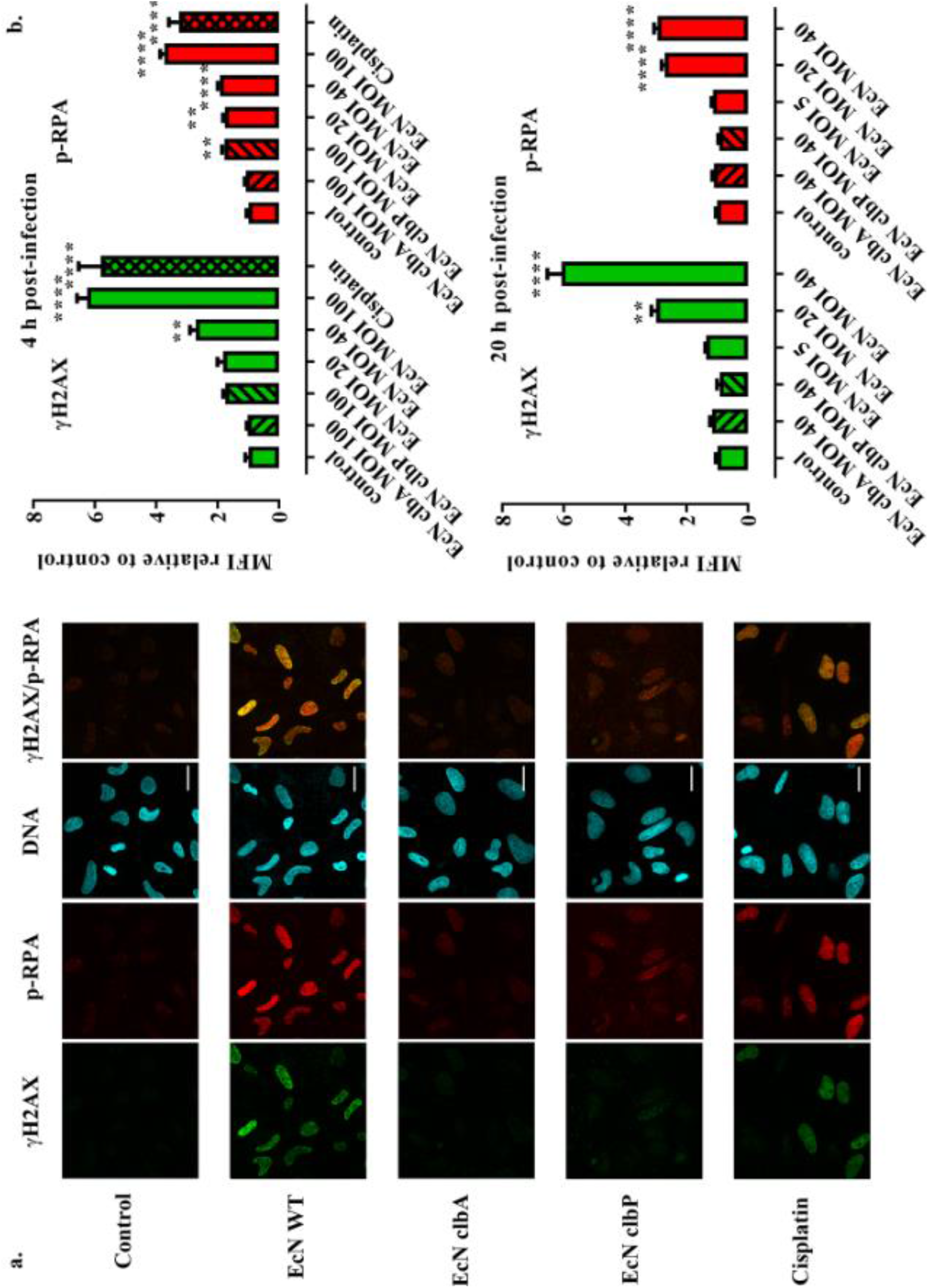
Formation of phosphorylated RPA and H2AX nuclear repair foci in HeLa cells infected with *E. coli* Nissle. (a) HeLa cells were exposed 4 hours to *E. coli* Nissle (EcN) or the *clbA* or *clbP* mutants (MOI = 100) or treated with cisplatin, then immunostained for phosphorylated H2AX (γH2AX) and phosphorylated RPA (p-RPA) 4 hours later. DNA was counterstained with DAPI. Bar = 20 μm. (b) Cells were infected with given MOI and immunostained at 4 and 20 hours after infection. The mean fluorescence intensity (MFI) of γH2AX and p-RPA within the nuclei, relative to that in control uninfected cells, was determined by image analysis using a macro in ImageJ. The means and standard errors, measured in at least 70 nuclei for each group, are shown. ** P<0.01, **** P<0.0001 (one-way ANOVA with Dunnett posttest, compared to control).

### Exposure to low numbers of Nissle 1917 induces abnormal mitosis and increased gene mutation frequency

Infection with colibactin-producing *E. coli* at low MOI can lead to incomplete DNA repair in a subset of the cell population, allowing cell division to restart and formation of aberrant anaphases, and ultimately increased gene mutation frequency (Cuevas-Ramos et al., 2010). We thus tested whether infection with Nissle 1917 induced these phenotypes, in epithelial CHO cells that have stable chromosomes and are amenable to gene mutation assay. CHO cells exposed to low numbers of wild-type Nissle 1917 showed abnormal mitotic figures 20 h after infection (Fig 3a). We observed lagging chromosomes, multipolar mitosis and anaphase DNA bridges in cells infected with Nissle 1917, or treated with cisplatin (Fig 3ab). The abnormal mitotic index increased with the MOI of the wild-type Nissle 1917 strain, whereas it remained at background level in cells exposed to the highest MOI of the *clbA* or *clbP* mutants (Fig. 3b).

**Figure 3.**
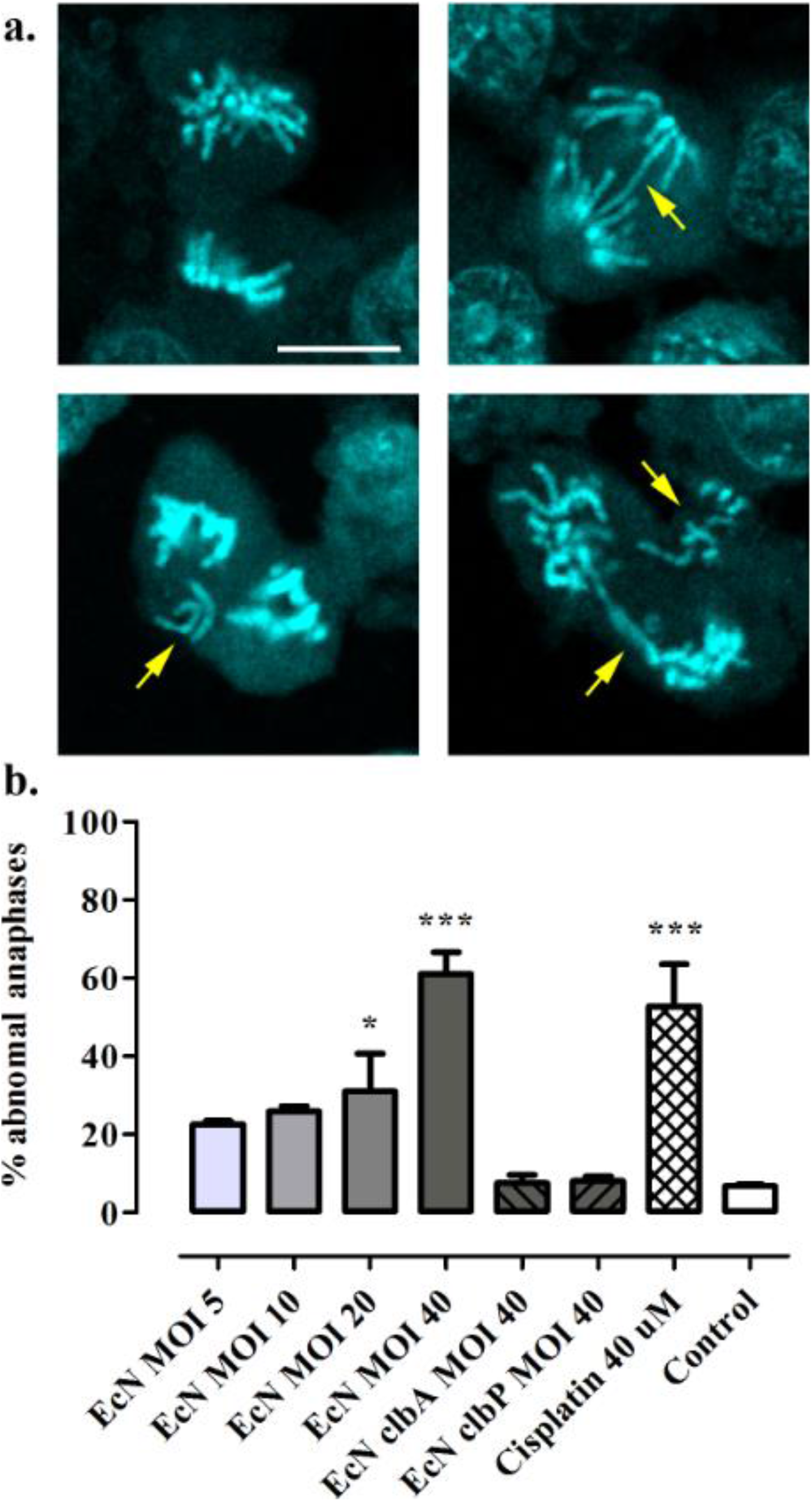
Infection with *E. coli* Nissle induces aberrant anaphases. (a) Anaphase bridges, lagging chromosomes and multipolar mitosis (arrows) in CHO cells 20 hours following infection with *E. coli* Nissle. DNA was stained with DAPI and observed by confocal microscopy. Bar = 20 μm. (b) Aberrant anaphase index in CHO cells 20 hours following infection with EcN at given MOI, or with the *clbA* and *clbP* mutants, or treatment with cisplatin. The means and standard errors, measured in three independent experiments, are shown. * P<0.05, ***P<0.001 (one-way ANOVA with Dunnett posttest compared to control).

Mitotic errors can lead to an accumulation of DNA damage, which in turn favors gene mutations (Chatterjee and Walker, 2017; Levine and Holland, 2018). We thus next assessed gene mutation frequencies at the hypoxanthine-guanine phosphoribosyltransferase (*hprt*) loci after infection of CHO cells (Table). We found a two-fold increase in 6-thioguanine-resistant (*hprt* mutant) colonies after infection with a MOI of 10 of the wild-type Nissle 1917 compared with uninfected cells or cells that were infected with the *clbA* or *clbP* mutant. The mutation frequency was similar to that previously observed with a laboratory *E. coli* strain hosting the *pks* island at the same MOI (Cuevas-Ramos et al., 2010), but did not reach statistical significance. Infection with a MOI of 20 Nissle 1917 resulted in a significant increase of *hprt* mutation frequency. Treatment with cisplatin also resulted in a significant increase of *hprt* mutants, with a mutation frequency similar to that reported in the literature (Silva et al., 2005). We conclude that Nissle 1917 is mutagenic.

### Nissle 1917 induces DNA damage to intestinal cells *in vivo*

To test whether Nissle 1917 produces colibactin *in vivo* in the gut lumen and induces DNA damages to intestinal cells, we first used a simplified model of intestinal colonization; adult axenic Balb/c mice were inoculated with Nissle 1917 or the *clbA* mutant, or with sterile PBS. Seven days after inoculation, the mice were sacrificed, fecal and intestinal tissue samples were collected. The mice mono-associated with Nissle 1917 or hosting the *clbA* mutant exhibited similar fecal counts of ~10^9^ CFU/g of feces. We assessed by immune-histology histone γH2AX in the colon. Nuclear γH2AX foci were readily observed in the enterocytes exposed to Nissle 1917, but not in animals inoculated with the *clbA* mutant, which exhibited background γH2AX levels similar to that of the axenic controls (Fig. 4ab).

**Figure 4.**
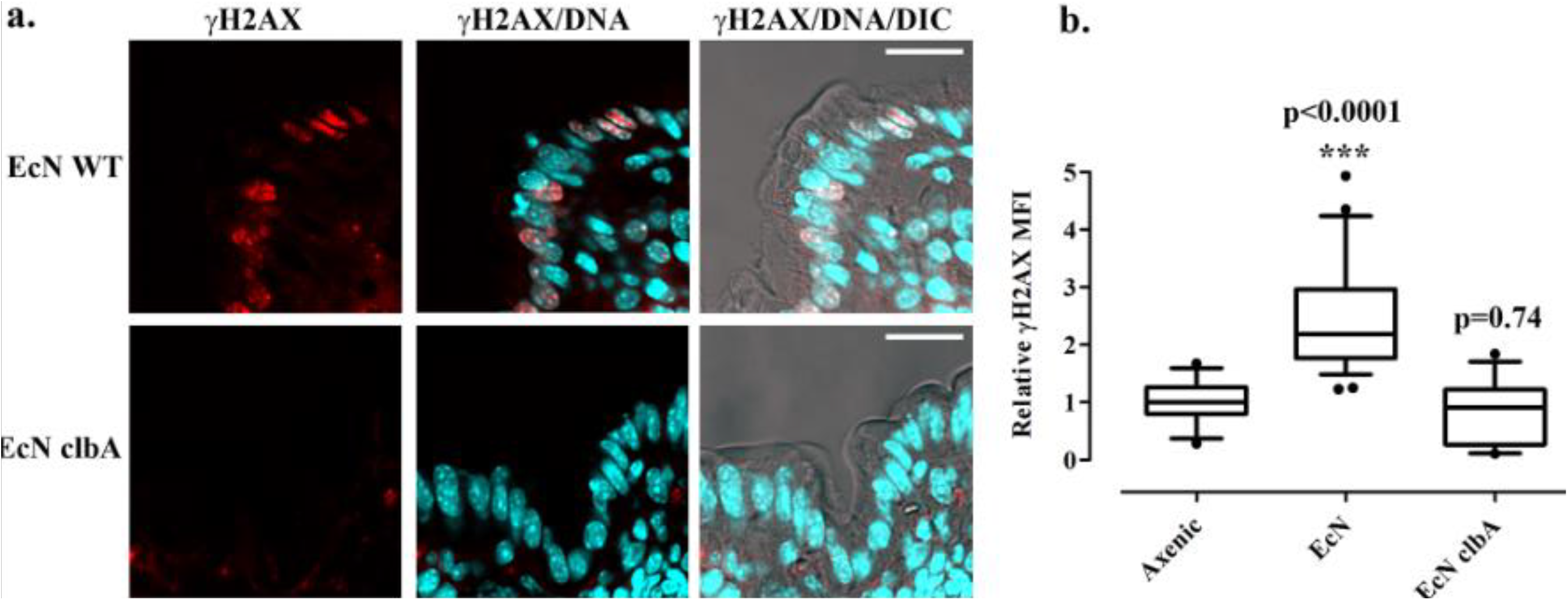
γH2AX foci in gut cells of mice mono-associated with *E. coli* Nissle 1917. Adult Balb/c mice were mono-colonized 7 days with wild-type *E. coli* Nissle 1917 (EcN WT) or the *clbA* mutant, or kept axenic. (a) γH2AX in histological sections of the colon was examined by immunofluorescence and confocal microscopy (red). DNA was counterstained with DAPI, and the tissue was visualized by differential interference contrast (DIC). Bars = 10 μm. (b) The mean fluorescence intensity (MFI) of γH2AX within the nuclei, relative to that measured in the axenic animals, was determined by automated image analysis in ImageJ. The whiskers show the median, 10-90 percentile and outliers, measured in at least 20 microscopic fields in 3 axenic and 5 mono-associated animals. The result of a one-way ANOVA with Dunnett posttest compared to axenic is shown.

Nissle 1917 is used not only in adults but also in infants and toddlers. To further examine production of colibactin *in vivo*, we used a second *in vivo* model in which 8-days old Swiss mouse pups were given *per os* ~10^8^ CFU of Nissle 1917 or the *clbP* mutant or PBS. Six hours after inoculation, the intestinal epithelium was examined for formation of γH2AX foci. Animals treated with Nissle 1917 exhibited significant levels of nuclear γH2AX compared to controls treated with PBS (Fig 5). In contrast, the animals treated with the *clbP* mutant that does not produce colibactin showed background levels of γH2AX (Fig 5). Together these results indicated that Nissle 1917 induces *in vivo* DNA damage to epithelial cells.

**Figure 5.**
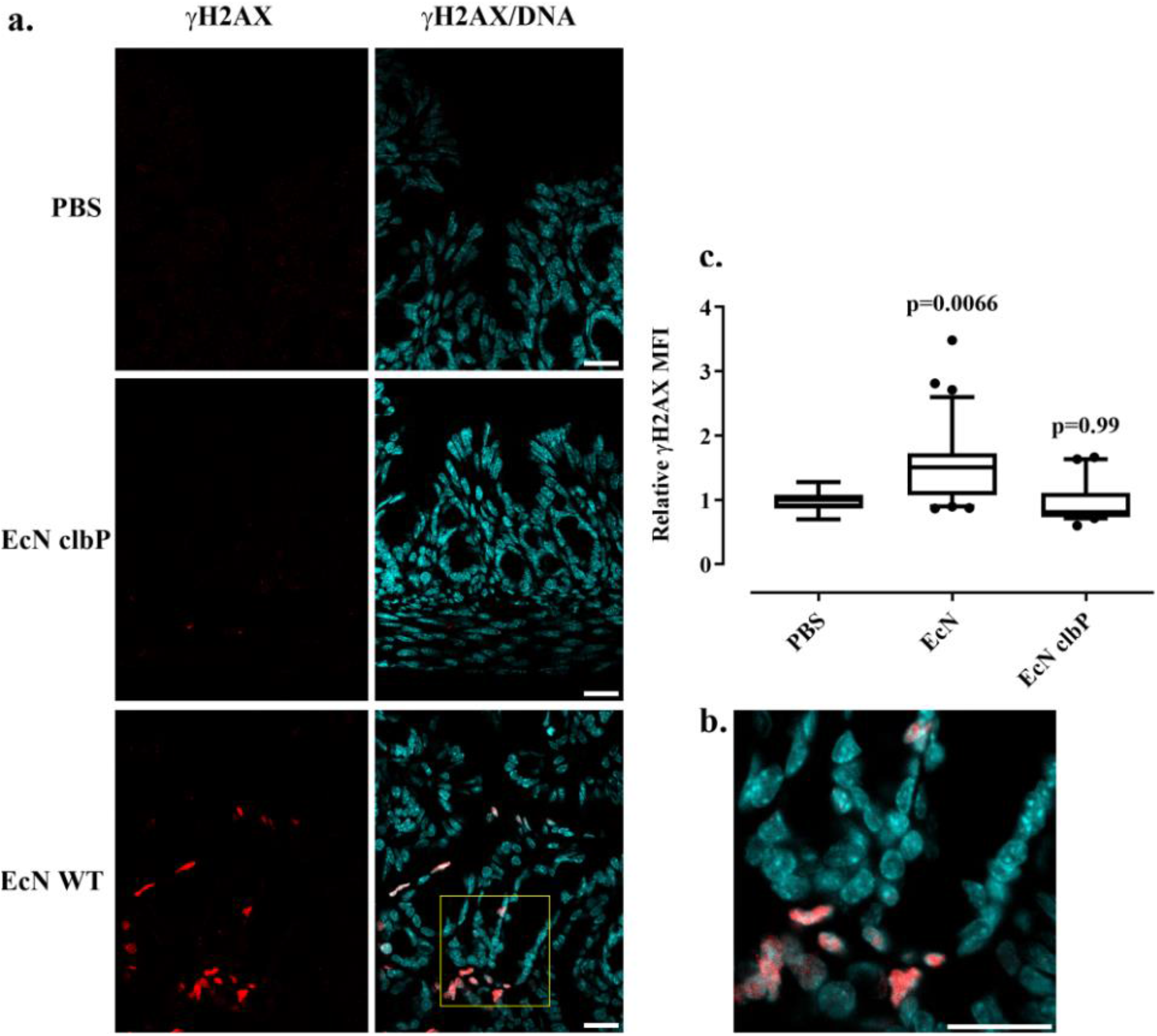
γH2AX foci in gut cells of mouse pups treated by *E. coli* Nissle 1917 by gavage. (a) Mice pups were given orally approximatively 2.5×10^8^ wild-type *E. coli* Nissle (EcN) or the *clbP* mutant, or the PBS vehicle, then sacrificed 6 hours later. Phosphorylated H2AX (γH2AX) in histological sections of the intestinal epithelium was examined by immunofluorescence (red) and confocal microscopy. DNA was counterstained with DAPI. Bars = 20 μm. (b) Close-up of the region shown in yellow. Bar = 20 μm. (c) The mean fluorescence intensity (MFI) of γH2AX within the nuclei, relative to that in the controls, was determined by automated image analysis in ImageJ. The whiskers show the median, 10-90 percentile and outliers, measured in at least 10 microscopic fields for each group in 3 controls (PBS) or 5 treated (Nissle 1917 or clbP mutant) animals. The result of a one-way ANOVA with Dunnett posttest compared to PBS is shown.

## Discussion

The identification of colibactin mutation signature in human colorectal cancer tissues (Dziubańska-Kusibab et al., 2020; Pleguezuelos-Manzano et al., 2020; Terlouw et al., 2020) and also in colonic crypts from healthy individuals under the age of ten (Lee-Six et al., 2019) proves that colibactin is expressed within the human gut (including in child), and links colibactin exposure to colorectal cancer. Colibactin is now a suspected prooncogenic driver especially in IBD patients (Dubinsky et al., 2020). Nissle 1917 has been used as a probiotic for various clinical applications since its isolation more than 100 years ago. It has shown some efficacy to treat IBDs such as Crohn’s disease and ulcerative colitis. In this study, we demonstrate that Nissle 1917 synthesizes colibactin, *in vitro* and *in vivo* in the gut lumen, and inflicts mutagenic DNA damages. Even in low numbers, DNA crosslinks are catastrophic damages that obstruct basic DNA processes, since they prevent the strand separation required for polymerase functions. The crosslinks notably perturb the replication machinery, resulting in replication stress, accumulation of DNA bound by RPA, activation of the kinase ATR that in turn phosphorylates RPA and histone variant H2AX (Bossuet-Greif et al., 2018; Maréchal and Zou, 2015; Vassin et al., 2009). We observed that cells exposed to Nissle 1917 at low MOI (hence numbers of bacteria more relevant to those occurring *in vivo*) entered an error-prone repair pathway, exhibiting mitotic aberrations and increased gene mutation frequency, similar to that observed with other *pks+ E. coli* strain (Cuevas-Ramos et al., 2010; Iftekhar et al., 2021). Thus, Nissle 1917 is genotoxigenic and mutagenic. This is of concern for patients and participants in clinical trials using Nissle 1917, such as the trial in Finland in which more than 250 young children will be inoculated with this strain (https://clinicaltrials.gov/ct2/show/NCT04608851)

Our results stand in contrast to that reported by Dubbert and colleagues who claimed that Nissle 1917 does not have detectable mutagenic activity using standard tests (Dubbert et al., 2020). However, the assays they used cannot detect colibactin-associated mutagenic damage. Indeed, to examine whether Nissle 1917 could induce mutagenic DNA damages, Dubbert used an Ames test in which *Salmonella typhimurium* reporter bacteria were exposed to Nissle 1917 and then *Salmonella* growth was expected upon mutagenesis. However, *Salmonella* bacteria are readily killed by the siderophores-microcins produced by Nissle 1917 (Massip et al., 2020; Sassone-Corsi et al., 2016) and thus the absence of growth of the reporter bacteria was incorrectly interpreted as an absence of effect of colibactin. In addition, Dubbert used a standard comet assay that can detect a variety of DNA lesions through electrophoresis of broken DNA, but which cannot detect DNA crosslinks that inhibit DNA electrophoretic mobility (Bossuet-Greif et al., 2018; Merk and Speit, 1999; Wilson et al., 2019). Thus, the standard assays used by Dubbert were inappropriate, in contrast to the assays used in the present and other works (Vizcaino and Crawford, 2015; Wilson et al., 2019), to highlight the DNA-damaging activity and genotoxicity of colibactin produced by Nissle 1917.

We demonstrate using two mouse models that Nissle 1917 synthesize colibactin in the gut and induces DNA damage in intestinal cells. Obviously, these mouse models do not fully recapitulate the human intestine, in particular its complex microbiota, epithelial and intestinal barrier functions. However, in human patients Nissle 1917 is typically used in the context of IBDs, where the gut is inflamed, the intestinal barrier is dysfunctional and the microbiota is dysbiotic. Importantly, intestinal inflammation was shown to upregulate *pks* genes (Arthur et al., 2014; Dubinsky et al., 2020; Yang et al., 2020). Inflammation and dysbiosis are also known to allow the expansion of the *E. coli* population, including that of Nissle 1917, alongside the epithelium (Cevallos et al., 2019; Dejea et al., 2018; Zhu et al., 2019). Moreover, Nissle 1917 is typically administered in very high numbers (2.5-25×10^9^ bacteria in adults, 10^8^ in infants), repeatedly (1-4 times daily), for weeks or even longer in case of ulcerative colitis. Nissle 1917 has been reported to persist in the human gut for months after inoculation (Lodinová-Zádniková and Sonnenborn, 1997). Thus, patients treated with this probiotic can be exposed chronically to high numbers of colibactin-producing bacteria, especially in an inflamed context that favor colibactin production, and consequently could be exposed to high levels of mutagenic colibactin. These conditions were shown to promote colon tumorigenesis in colorectal cancer (Arthur et al., 2012).

Nissle 1917 is an increasingly popular choice to engineer live biotherapeutics (i.e. bacteria genetically designed to treat or prevent a disease) (Charbonneau et al., 2020). For example, Nissle 1917 has been used successfully as a chassis to deliver an anti-biofilm enzyme against *P. aeruginosa* (Hwang et al., 2017), or a microcin induced upon sensing of *Salmonella* infection (Palmer et al., 2018). Engineered strains of Nissle 1917 have also been constructed to treat obesity through production of N- acylphosphatidylethanolamine (Chen et al., 2014) or to express a phenylalanine-metabolizing enzyme in response to the anoxic conditions in the gut, to treat phenylketonuria (Isabella et al., 2018). Considering the widespread use of Nissle 1917, as a probiotic and as a platform to develop live bacterial therapeutics, ensuring its safety is of paramount importance. Genotoxic carcinogens are classically conceived to represent a risk factor with no threshold dose, because little numbers or even one DNA lesion may result in mutation and increased tumor risk (Hartwig et al., 2020). Production of mutagenic colibactin by Nissle 1917 is thus a serious health concern that must be addressed.

## Methods

### *E. coli* EcN strain, mutants and culture

The *E. coli* strain Nissle 1917 used in this study was obtained from Dr. Ulrich Dobrindt (University of Münster). The *clbA* and *clbP* isogenic mutants were described previously (ref Ollier et Nat Comm). Before infection, the bacteria were grown overnight at 37°C with 240 RPM agitation in 5 mL of Lennox L broth (LB, Invitrogen) then diluted 1/20 in pre-warmed DMEM 25 mM Hepes (Invitrogen) and incubated at 37°C with 240 RPM agitation to reach exponential phase (OD600=0.4 to 0.5).

### *In vitro* DNA crosslinking assay

3×10^6^ bacteria or numbers given in the text were inoculated in 100 μl of DMEM 25 mM Hepes, incubated at 37°C for 3.5 hours, then EDTA (1 mM) and 400 ng of linearized (BamHI) pUC19 DNA were added and further incubated 40 minutes. As controls, DNA was left uninfected or was treated with 100 or 200 μM cisplatin (Sigma). Following a centrifugation to pellet the bacteria, the DNA was purified using Qiagen PCR DNA purification kit before analysis by denaturing gel electrophoresis.

### Denaturing gel DNA electrophoresis

1% agarose gels prepared in a 100 mM NaCl 2 mM EDTA pH 8 solution were soaked 16 hours in 40 mM NaOH 1 mM EDTA electrophoresis running buffer. DNA electrophoresis was performed at room temperature, 45 min at 1 V/cm then 2 h at 2 V/cm. Following neutralization by serial washes in 150 mM NaCl 100 mM Tris pH 7.4, DNA was stained with Gel Red (Biotium) and photographed with flat-field correction and avoiding CCD pixel saturation in a Biorad Chemidoc XRS system. Images were analyzed using NIH ImageJ: the background was subtracted (100 pixels rolling ball) then the lane profiles were plotted and the area of DNA peaks were measured.

### Cell culture and infection

HeLa and CHO cells were cultivated in a 37°C 5% CO_2_ incubator and maintained by serial passage in DMEM Glutamax or MEMα (Invitrogen) respectively, both supplemented with 10% fetal calf serum (FCS), 50 μg/ml gentamicin and 1% non-essential amino acids (Invitrogen). 3×10^5^ cells/well were seeded in 6-wells plates (TPP) or 3.5×10^4^ cells/well in 8-chambers slides (Falcon) and grown 24 hours. Cells were washed 3 times in HBSS (Invitrogen) before infection in DMEM 25 mM HEPES at given multiplicity of infection (MOI = number of bacteria per cell at the onset of infection). Following the 4 hours co-culture, the cells were washed 3 times with HBSS then incubated in complete cell culture medium supplemented with 200 μg/ml gentamicin for the indicated times (0, 4 or 20 hours) before analysis.

### *In cellulo* genomic DNA crosslinking assay

The cells were infected 4 hours or treated 4 hours with 100 μM cisplatin (Sigma), then collected immediately by trypsination. The cell genomic DNA was purified with Qiagen DNeasy Blood and Tissue kit and analyzed by denaturing gel electrophoresis.

### Abnormal anaphase scoring

Abnormal anaphase quantification was done as described (Luo et al., 2004). Briefly, 3 hours after the end of infection, the cells were trapped in premetaphase by treatment with 0.6 μg/ml nocodazole and released 55 min without nocodazole to reach anaphase. The slides were fixed, stained with DAPI and examined by confocal microscopy as described below. The anaphases were scored in three independent experiments.

### Gene mutation assay

CHO cells were treated 4 days with culture medium supplemented with 10mM deoxycytidine 200mM hypoxanthine 0.2mM aminoprotein and 17.5mM thymidine (Sigma) to eliminate preexisting *hprt* mutants. CHO were infected 4 hours with Nissle 1917 or *clbA* or *clbP* mutants, or treated with cisplatin, then washed and cultured one week in normal cell culture medium and passaged in 10 cm dishes seeded with 3×10^5^ cells using culture medium supplemented with 30 μM 6-thioguanine (6-TG, Sigma). Cells were also plated without 6-TG to determine plating efficiency. The culture media was changed twice a week for 21 days. Then plates were fixed with 4% formaldehyde and stained with methylene blue.

### Animal studies

All procedures were carried out according to European and French guidelines for the care and use of laboratory animals. The experimentations were approved by Regional Council of Ethics for animal experimentation. SPF pregnant Swiss mice obtained from Janvier (Le Genest, St Isle, France) were housed under SPF conditions in the Inserm Purpan animal facility (Toulouse, France). Eight days old mice pups received *per os* a drop (approximately 25 μl) of bacteria suspended (10^10^ CFU/ml) in PBS, and were sacrificed 6 hours later (protocols 16-U1220-JPN/FT-010 and 17-U1220-EO/PM-461). Germ-free Balb/c mice were housed in the breeding facility of ANAXEM (INRAE, UMR1319 MICALIS, Jouy-en- Josas, France). Axenic animals were inoculated once by intragastric gavage with 10^8^ bacteria suspended in PBS, and sacrificed 7 days later (protocol APAFIS#3441-2016010614307552 v1). Intestinal tissue samples were fixed 24 hours in neutral buffered formalin, dehydrated in ethanol and embedded in paraffin.

### γH2AX and p-RPA immunofluorescence analysis

4 or 20 hours after infection, HeLa cells were pre-extracted 5 min in PBS 0.1% Triton X-100 before a 30 min fixation in PBS 4% formaldehyde. Following permeabilization in 0.1% Triton X-100 and blocking in MAXblock medium (Active Motif), the cells were stained 3 hours with antibodies against γH2AX (1:500, JBW301, Millipore) and S33p-RPA32 (1:500, A300-264A, Bethyl) diluted in MAXblock 0.05% Triton X-100. The cells were washed 3 times in PBS 0.05% Triton X-100 and incubated 1 h with anti-mouse AlexaFluor 488 and anti-rabbit AlexaFluor 568 (Invitrogen) diluted 1:500 in MAXblock medium with 1 μg/ml DAPI (Sigma). The cells were washed again, mounted in Fluoroshield medium (Sigma), and examined with a Leica SP8 laser scanning confocal microscope in sequential mode. The mean fluorescence intensities (MFI) of γH2AX and p-RPA within the nuclei were analyzed using a NIH ImageJ macro: the nuclei were identified in the DNA image (following a 0.5 μm Gaussian blur and default auto-threshold) and copied in the ROI manager to measure their corresponding MFI in the green and red channels.

For immunohistological staining of γH2AX in intestinal tissues, sections (5 or 8 μm) were deparaffinized by serial washes in xylene and ethanol, then rehydrated with water. The antigens were unmasked in HBSS 0.05% trypsin 0.02% EDTA at 37°C for 6 min then in sodium citrate buffer (10 mM sodium citrate, 0.05% Tween 20, pH 6.0) for 30 min at 80-95°C. Following a 1 hour cooling to room temperature and blocking 1 hour in 0.3% Triton X-100 MAXblock medium, the tissues were stained 16 hours at 4°C with primary antibodies against γH2AX (1:200, 20E3, Cell Signaling Technology) diluted in the blocking medium. The slides were washed 3 times in PBS 0.05% Triton X-100 and incubated 1 h with anti-rabbit AlexaFluor 568 diluted 1:200 in MAXblock medium with 1 μg/ml DAPI. The slides were washed again, mounted and examined as above.

**Table 1:** *hprt* mutant frequencies (MF) following infection with *E. coli* Nissle 1917 at given multiplicity of infection (MOI), or with the *clbA* or *clbP* mutants, or 1 h treatment with cisplatin. The values are the mean and standard error of three independent infection experiments. Statistical analysis compared to control was performed using a two tailed t-test on the log transformed data.

| **Treatment** | **MF x10<sup>-6</sup>** | **p** |
| --- | --- | --- |
| Control | 5.99 ± 0.98 | - |
| Cisplatin 10 μM | 25.25 ± 5.83 | ** 0.006 |
| Cisplatin 15 μM | 47.62 ± 12.60 | ** 0.002 |
| EcN MOI 5 | 5.66 ± 0.71 | 0.685 |
| EcN MOI 10 | 11.98 ± 5.99 | 0.425 |
| EcN MOI 20 | 14.49 ± 8.37 | * 0.023 |
| EcN *clbA* MOI 20 | 5.46 ± 0.28 | 0.450 |
| EcN *clbP* MOI 20 | 4.94 ± 0.51 | 0.168 |

## Statistical analyses

Statistical analyses were performed using Graph Pad Prism 9. Analysis of mutant frequencies was performed using a two tailed t-test on the log transformed data, to ensure data normality and to correct variance heterogeneity (Silva et al., 2005).

## Acknowledgements

We thank Sophie Allart for technical assistance at the cellular imaging facility of Inserm UMR 1291, Toulouse. This work was funded by a French governmental grant from the Institut National Du Cancer (INCA PLBIO13-123). CC was the recipient of a scholarship (“poste d’accueil”) from Inserm. JPM was funded by a fellowship (“AgreenSkills+”) from the EU’s Seventh Framework Program FP7-609398. The funders had no role in study design, data collection and interpretation, or the decision to submit the work for publication.

---

**Supplementary figure 1.**
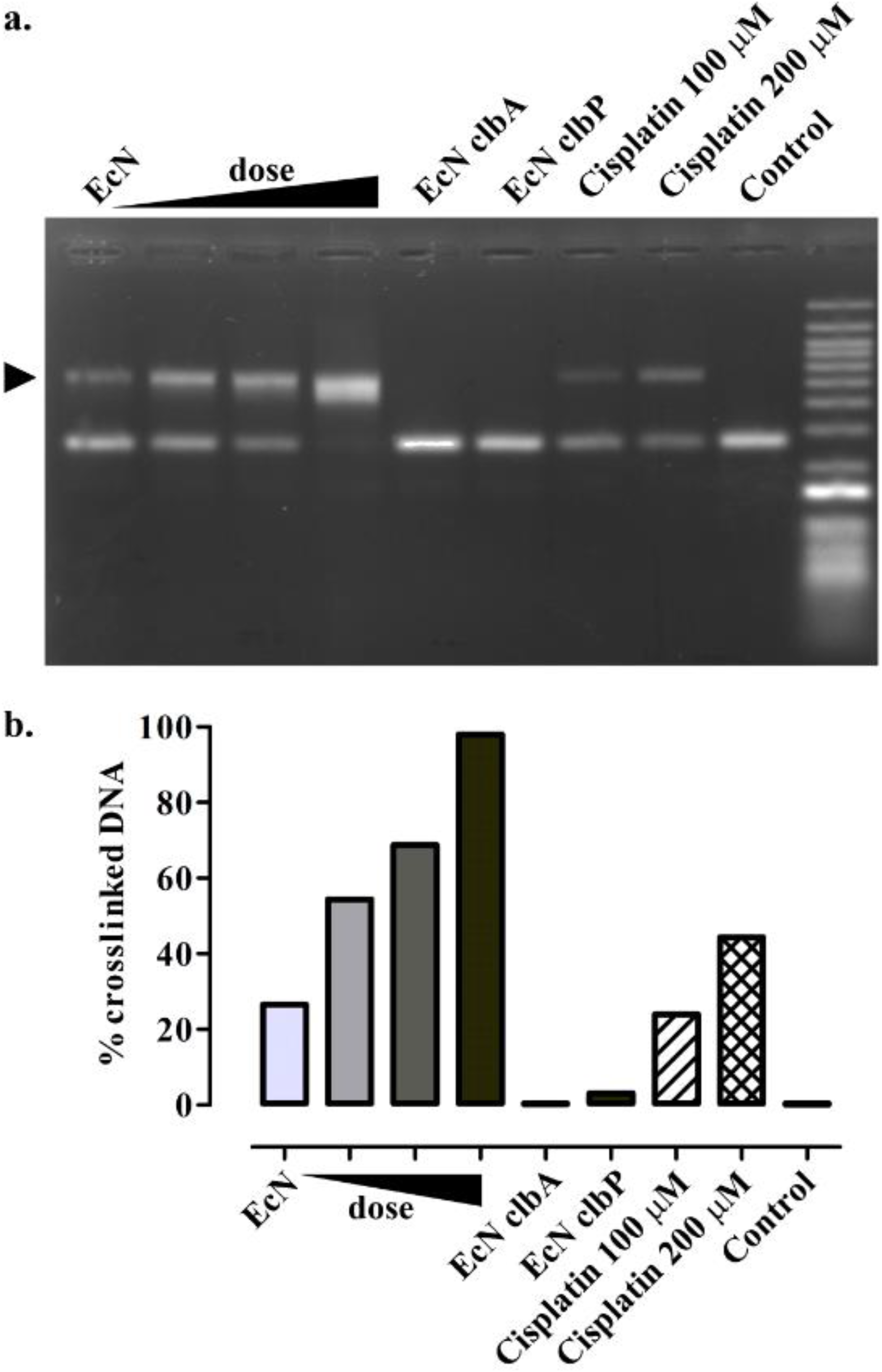
*E. coli* Nissle induces interstrand crosslinks in exogenous DNA. (a) Linearized plasmid double-strand DNA was incubated 40 minutes with *E. coli* Nissle (EcN) (inoculum of 0.75, 1.5, 3, or 6×10^6^ bacteria in 100 μl grown 3.5 hours) or with the *clbA* or *clbP* mutants (6×10^6^ bacteria in 100 μl) or treated 4 h with cisplatin, and then analyzed by denaturing gel electrophoresis. (b) Quantification of panel a; the percentages of the DNA signal in the upper cross-linked band relative to the total DNA signal in the lane was determined by image analysis.

## References

Arthur, J.C., Perez-Chanona, E., Mühlbauer, M., Tomkovich, S., Uronis, J.M., Fan, T.-J., Campbell, B.J., Abujamel, T., Dogan, B., Rogers, A.B., et al. (2012). Intestinal Inflammation Targets Cancer-Inducing Activity of the Microbiota. Science.

Arthur, J.C., Gharaibeh, R.Z., Mühlbauer, M., Perez-Chanona, E., Uronis, J.M., McCafferty, J., Fodor, A.A., and Jobin, C. (2014). Microbial genomic analysis reveals the essential role of inflammation in bacteria-induced colorectal cancer. Nat Commun 5, 4724.

Bossuet-Greif, N., Vignard, J., Taieb, F., Mirey, G., Dubois, D., Petit, C., Oswald, E., and Nougayrède, J.-P. (2018). The Colibactin Genotoxin Generates DNA Interstrand Cross-Links in Infected Cells. MBio 9, e02393–17.

Brotherton, C.A., and Balskus, E.P. (2013). A prodrug resistance mechanism is involved in colibactin biosynthesis and cytotoxicity. J. Am. Chem. Soc. 135, 3359–3362.

Cevallos, S.A., Lee, J.-Y., Tiffany, C.R., Byndloss, A.J., Johnston, L., Byndloss, M.X., and Bäumler, A.J. (2019). Increased Epithelial Oxygenation Links Colitis to an Expansion of Tumorigenic Bacteria. MBio 10.

Charbonneau, M.R., Isabella, V.M., Li, N., and Kurtz, C.B. (2020). Developing a new class of engineered live bacterial therapeutics to treat human diseases. Nature Communications 11, 1738.

Chatterjee, N., and Walker, G.C. (2017). Mechanisms of DNA damage, repair, and mutagenesis. Environmental and Molecular Mutagenesis 58, 235–263.

Chen, Z., Guo, L., Zhang, Y., Walzem, R.L., Pendergast, J.S., Printz, R.L., Morris, L.C., Matafonova, E., Stien, X., Kang, L., et al. (2014). Incorporation of therapeutically modified bacteria into gut microbiota inhibits obesity. J Clin Invest 124, 3391–3406.

Cougnoux, A., Dalmasso, G., Martinez, R., Buc, E., Delmas, J., Gibold, L., Sauvanet, P., Darcha, C., Déchelotte, P., Bonnet, M., et al. (2014). Bacterial genotoxin colibactin promotes colon tumour growth by inducing a senescence-associated secretory phenotype. Gut gutjnl-2013-305257.

Cuevas-Ramos, G., Petit, C.R., Marcq, I., Boury, M., Oswald, E., and Nougayrède, J.-P. (2010). Escherichia coli induces DNA damage in vivo and triggers genomic instability in mammalian cells. Proc. Natl. Acad. Sci. U.S.A. 107, 11537–11542.

Dejea, C.M., Fathi, P., Craig, J.M., Boleij, A., Taddese, R., Geis, A.L., Wu, X., DeStefano Shields, C.E., Hechenbleikner, E.M., Huso, D.L., et al. (2018). Patients with familial adenomatous polyposis harbor colonic biofilms containing tumorigenic bacteria. Science 359, 592–597.

Dubbert, S., Klinkert, B., Schimiczek, M., Wassenaar, T.M., and von Bünau, R. (2020). No Genotoxicity Is Detectable for Escherichia coli Strain Nissle 1917 by Standard In Vitro and In Vivo Tests. Eur J Microbiol Immunol (Bp) 10, 11–19.

Dubinsky, V., Dotan, I., and Gophna, U. (2020). Carriage of Colibactin-producing Bacteria and Colorectal Cancer Risk. Trends in Microbiology 28, 874–876.

Dziubańska-Kusibab, P.J., Berger, H., Battistini, F., Bouwman, B.A.M., Iftekhar, A., Katainen, R., Cajuso, T., Crosetto, N., Orozco, M., Aaltonen, L.A., et al. (2020). Colibactin DNA-damage signature indicates mutational impact in colorectal cancer. Nat. Med.

Floch, M.H., Walker, W.A., Madsen, K., Sanders, M.E., Macfarlane, G.T., Flint, H.J., Dieleman, L.A., Ringel, Y., Guandalini, S., Kelly, C.P., et al. (2011). Recommendations for probiotic use-2011 update. J Clin Gastroenterol 45 Suppl, S168–171.

Hartwig, A., Arand, M., Epe, B., Guth, S., Jahnke, G., Lampen, A., Martus, H.-J., Monien, B., Rietjens, I.M.C.M., Schmitz-Spanke, S., et al. (2020). Mode of action-based risk assessment of genotoxic carcinogens. Arch Toxicol 94, 1787–1877.

Homburg, S., Oswald, E., Hacker, J., and Dobrindt, U. (2007). Expression analysis of the colibactin gene cluster coding for a novel polyketide in Escherichia coli. FEMS Microbiol. Lett. 275, 255–262.

Hwang, I.Y., Koh, E., Wong, A., March, J.C., Bentley, W.E., Lee, Y.S., and Chang, M.W. (2017). Engineered probiotic Escherichia coli can eliminate and prevent Pseudomonas aeruginosa gut infection in animal models. Nature Communications 8, 15028.

Iftekhar, A., Berger, H., Bouznad, N., Heuberger, J., Boccellato, F., Dobrindt, U., Hermeking, H., Sigal, M., and Meyer, T.F. (2021). Genomic aberrations after short-term exposure to colibactin-producing E. coli transform primary colon epithelial cells. Nat Commun 12, 1003.

Isabella, V.M., Ha, B.N., Castillo, M.J., Lubkowicz, D.J., Rowe, S.E., Millet, Y.A., Anderson, C.L., Li, N., Fisher, A.B., West, K.A., et al. (2018). Development of a synthetic live bacterial therapeutic for the human metabolic disease phenylketonuria. Nat Biotechnol 36, 857–864.

Kruis, W., Frič, P., Pokrotnieks, J., Lukáš, M., Fixa, B., Kaščák, M., Kamm, M.A., Weismueller, J., Beglinger, C., Stolte, M., et al. (2004). Maintaining remission of ulcerative colitis with the probiotic Escherichia coli Nissle 1917 is as effective as with standard mesalazine. Gut 53, 1617–1623.

Lee-Six, H., Olafsson, S., Ellis, P., Osborne, R.J., Sanders, M.A., Moore, L., Georgakopoulos, N., Torrente, F., Noorani, A., Goddard, M., et al. (2019). The landscape of somatic mutation in normal colorectal epithelial cells. Nature 574, 532–537.

Levine, M.S., and Holland, A.J. (2018). The impact of mitotic errors on cell proliferation and tumorigenesis. Genes Dev. 32, 620–638.

Lodinová-Zádniková, R., and Sonnenborn, U. (1997). Effect of preventive administration of a nonpathogenic Escherichia coli strain on the colonization of the intestine with microbial pathogens in newborn infants. Biol Neonate 71, 224–232.

Luo, L.Z., Werner, K.M., Gollin, S.M., and Saunders, W.S. (2004). Cigarette smoke induces anaphase bridges and genomic imbalances in normal cells. Mutation Research/Fundamental and Molecular Mechanisms of Mutagenesis 554, 375–385.

Marcq, I., Martin, P., Payros, D., Cuevas-Ramos, G., Boury, M., Watrin, C., Nougayrède, J.-P., Olier, M., and Oswald, E. (2014). The genotoxin colibactin exacerbates lymphopenia and decreases survival rate in mice infected with septicemic Escherichia coli. J. Infect. Dis. 210, 285–294.

Maréchal, A., and Zou, L. (2015). RPA-coated single-stranded DNA as a platform for post-translational modifications in the DNA damage response. Cell Research 25, 9–23.

Martin, P., Marcq, I., Magistro, G., Penary, M., Garcie, C., Payros, D., Boury, M., Olier, M., Nougayrède, J.-P., Audebert, M., et al. (2013). Interplay between siderophores and colibactin genotoxin biosynthetic pathways in Escherichia coli. PLoS Pathog. 9, e1003437.

Massip, C., Chagneau, C.V., Boury, M., and Oswald, E. (2020). The synergistic triad between microcin, colibactin, and salmochelin gene clusters in uropathogenic Escherichia coli. Microbes Infect. 22, 144–147.

McCarthy, A.J., Martin, P., Cloup, E., Stabler, R.A., Oswald, E., and Taylor, P.W. (2015). The Genotoxin Colibactin Is a Determinant of Virulence in Escherichia coli K1 Experimental Neonatal Systemic Infection. Infect. Immun. 83, 3704–3711.

Merk, O., and Speit, G. (1999). Detection of crosslinks with the comet assay in relationship to genotoxicity and cytotoxicity. Environmental and Molecular Mutagenesis 33, 167–172.

Nissle, A. (1959). [On coli antagonism, dysbacteria and coli therapy]. Med Monatsschr 13, 489–491.

Nougayrède, J.-P., Homburg, S., Taieb, F., Boury, M., Brzuszkiewicz, E., Gottschalk, G., Buchrieser, C., Hacker, J., Dobrindt, U., and Oswald, E. (2006). Escherichia coli induces DNA double-strand breaks in eukaryotic cells. Science 313, 848–851.

Ou, B., Yang, Y., Tham, W.L., Chen, L., Guo, J., and Zhu, G. (2016). Genetic engineering of probiotic Escherichia coli Nissle 1917 for clinical application. Appl Microbiol Biotechnol 100, 8693–8699.

Palmer, J.D., Piattelli, E., McCormick, B.A., Silby, M.W., Brigham, C.J., and Bucci, V. (2018). Engineered Probiotic for the Inhibition of Salmonella via Tetrathionate-Induced Production of Microcin H47. ACS Infect Dis 4, 39–45.

Pleguezuelos-Manzano, C., Puschhof, J., Rosendahl Huber, A., van Hoeck, A., Wood, H.M., Nomburg, J., Gurjao, C., Manders, F., Dalmasso, G., Stege, P.B., et al. (2020). Mutational signature in colorectal cancer caused by genotoxic pks+ E. coli. Nature 580, 269–273.

Pradhan, S., and Weiss, A.A. (2020). Probiotic Properties of Escherichia coli Nissle in Human Intestinal Organoids. MBio 11.

Sassone-Corsi, M., Nuccio, S.-P., Liu, H., Hernandez, D., Vu, C.T., Takahashi, A.A., Edwards, R.A., and Raffatellu, M. (2016). Microcins mediate competition among Enterobacteriaceae in the inflamed gut. Nature 540, 280–283.

Silva, M.J., Costa, P., Dias, A., Valente, M., Louro, H., and Boavida, M.G. (2005). Comparative analysis of the mutagenic activity of oxaliplatin and cisplatin in the Hprt gene of CHO cells. Environmental and Molecular Mutagenesis 46, 104–115.

Terlouw, D., Suerink, M., Boot, A., Wezel, T. van, Nielsen, M., and Morreau, H. (2020). Recurrent APC Splice Variant c.835-8A>G in Patients With Unexplained Colorectal Polyposis Fulfilling the Colibactin Mutational Signature. Gastroenterology 159, 1612–1614.e5.

Vassin, V.M., Anantha, R.W., Sokolova, E., Kanner, S., and Borowiec, J.A. (2009). Human RPA phosphorylation by ATR stimulates DNA synthesis and prevents ssDNA accumulation during DNA-replication stress. J. Cell. Sci. 122, 4070–4080.

Vizcaino, M.I., and Crawford, J.M. (2015). The colibactin warhead crosslinks DNA. Nat Chem 7, 411–417.

Wilson, M.R., Jiang, Y., Villalta, P.W., Stornetta, A., Boudreau, P.D., Carrá, A., Brennan, C.A., Chun, E., Ngo, L., Samson, L.D., et al. (2019). The human gut bacterial genotoxin colibactin alkylates DNA. Science 363.

Yang, Y., Gharaibeh, R.Z., Newsome, R.C., and Jobin, C. (2020). Amending microbiota by targeting intestinal inflammation with TNF blockade attenuates development of colorectal cancer. Nature Cancer 1, 723–734.

Zhu, W., Miyata, N., Winter, M.G., Arenales, A., Hughes, E.R., Spiga, L., Kim, J., Sifuentes-Dominguez, L., Starokadomskyy, P., Gopal, P., et al. (2019). Editing of the gut microbiota reduces carcinogenesis in mouse models of colitis-associated colorectal cancer. Journal of Experimental Medicine 216, 2378–2393.

